# Advances in two photon scanning and scanless microscopy technologies for functional neural circuit imaging

**DOI:** 10.1101/036632

**Authors:** Simon R. Schultz, Caroline S. Copeland, Amanda J. Foust, Peter Quicke, Renaud Schuck

## Abstract

Recent years have seen substantial developments in technology for imaging neural circuits, raising the prospect of large scale imaging studies of neural populations involved in information processing, with the potential to lead to step changes in our understanding of brain function and dysfunction. In this article we will review some key recent advances: improved fluorophores for single cell resolution functional neuroimaging using a two photon microscope; improved approaches to the problem of *scanning* active circuits; and the prospect of *scanless* microscopes which overcome some of the bandwidth limitations of current imaging techniques. These advances in technology for experimental neuroscience have in themselves led to technical challenges, such as the need for the development of novel signal processing and data analysis tools in order to make the most of the new experimental tools. We review recent work in some active topics, such as region of interest segmentation algorithms capable of demixing overlapping signals, and new highly accurate algorithms for calcium transient detection. These advances motivate the development of new data analysis tools capable of dealing with spatial or spatiotemporal patterns of neural activity, that scale well with pattern size.

## 1 Two photon imaging of neural activity patterns

Understanding the principles by which neural circuits process information is one of the central problems of modern neuroscience. It is also one of the most important, as its resolution underpins our understanding of both natural brain function, and pathological neural processes. In order to make substantial advances in the treatment of disorders of neural circuits-from developmental disorders such as autistic spectrum disorders to neurodegenerative disorders such as Alzheimer’s Disease-it is necessary to improve our understanding of how such circuits process information.

A necessary step in reverse engineering the functional principles of neural circuits is to simultaneously observe the activity of local circuit elements-down to the resolution of individual neuronal cell bodies, and potentially even of subcellular elements such as dendrites. We refer to such observation as functional cellular neuroimaging. Spatially dense sampling of neurons is further motivated by the highly nonrandom connectivity between them-nearby neurons are much more likely to be connected than distal neurons [1], and thus more likely to be involved in common information processing circuitry. There are currently only two technologies that offer the prospect of dense recording from many neurons located within a local volume of tissue: one electrophysiological, one optical. The electrophysiological approach is polytrode recording, using silicon planar arrays of densely spaced electrodes [2] for extracellular recording. Multi-electrode array electrophysiological recording technology offers greater spike timing precision and accuracy at detection of individual action potentials than current optical approaches. However, use of this technology for spatial localisation and cell type classification of recorded cells is still a work in progress [3]. A further limitation is tissue damage and neuroinflammation from large penetrating electrodes.

In comparison, optical approaches to monitoring neural activity are relatively noninvasive, allow the precise spatial location of activity-related signals to be identified, and can be used in conjunction with genetically targeted co-labelling to aid cell type classification. The development of two photon microscopy in the early 1990s [4] allowed optical imaging with a very small point spread function (of a volume of the order of the wavelength cubed), meaning that it is possible to know precisely which neural structure generated an observed photon, in real time. In addition, two photon microscopy has advantages over single photon fluorescence imaging in terms of imaging depth (as longer wavelength light is used for multiphoton than single photon excitation of a given fluorophore), with depths of up to 1 mm having been achieved for imaging labelled neural structures in cerebral cortex [5], and depths of several mm achievable for imaging vasculature [6]. Three photon imaging appears likely to yield even deeper imaging of functional neural signals [7].

Two photon microscopy thus has great potential as a tool for reverse engineering the functional architecture of neural circuits. However, there are limitations: the relatively slow kinetics and/or limited amplitude of the fluorescence signal from many activity dependent fluorophores mean that the temporal resolution with which spike trains can currently be resolved is far inferior to electrophysiology. In addition, two photon laser scanning microscopy is essentially a point measurement technique (with images being built up by scanning). This means that there is a trade-off between sampling rate and the number of points being acquired. Finally, the development of signal processing and data analysis tools for this type of data is in a relative stage of infancy. All of these issues are, however, the object of much investigation, and the field is progressing rapidly. In this article we will review recent advances in activity dependent fluorophores, scanning algorithms for observing neural circuit activity, scanless approaches to imaging neural circuits, and recent developments in data analysis algorithms for two photon imaging.

## 2 Labelling neural circuits with activity-dependent fluorophores

Fluorescent indicators of neuronal activity fall into two major classes, those that detect changes in intracellular calcium concentration, and those sensitive towards membrane voltage. Within these two major classes, both synthetic indicators and genetically encoded proteins have been designed and implemented; these shall be discussed below. Other approaches, such as the use of sensors for sodium or potassium ions, or pH, are in principle possible, but are currently many orders of magnitude off the sensitivity required for use at cellular resolution in the study of intact neural networks. Current sensors for membrane potential imaging yield relatively small signals in response to a single action potential, due to both sensor signal to noise ratio (SNR) and the short duration of the action potential-although recent developments are promising (see Section 2.3).

For this reason, most work on imaging neural circuit activity to date has employed calcium sensors. Upon action potential initiation and propagation, intracellular neuronal calcium concentration can rise ten-to a hundred-times higher than under conditions of rest [8]. Whilst there are multiple sources from which this calcium is derived via a host of different mechanisms [for review see 9], voltage-gated calcium channels facilitating calcium influx from the extracellular space are the primary determinants of changes in intracellular calcium concentration.

### 2.1 *Synthetic calcium dyes*

The examination of neuronal activity via intracellular calcium levels was first enabled by bioluminescent calcium-binding photoproteins, such as aequorin [10, 11], and calcium-sensitive absorbance dyes, such as arsenazo III [12]. Whilst these tools provided the first insights into intracellular calcium dynamics [13–15], their limited application, in that they required loading on a single cell basis via micropipette due to cell membrane impermeability [16], meant that they were soon superseded.

The second generation of calcium-sensitive indicators comprised a family of buffers developed by hybridization of a fluorescent chromophore with a calcium-selective chelator, such as BAPTA orEGTA [17]. These dyes, which include quin-2, fura-2, indo-1, and fluo-3, could be loaded into tissue more easily via bolus loading than the single-cell approach required by aequorin or arsenazo III. Multi-cell bolus loading [18], essentially the pressure injection of a bolus or “ball” of dye directly into tissue, has proved to be an effective way to label large groups of cells. Of these dyes, fura-2 soon became a popular choice amongst neuroscientists due to its large changes in fluorescence [19], and its ability to enable quantitative determination of the calcium concentration [20, 21]. Upon binding of calcium ions, fura-2 undergoes intramolecular conformational changes that lead to a change in the emitted fluorescence. Interestingly, this change manifests as a decrease in fluorescence upon an increase in intracellular calcium levels, under two-photon imaging conditions [22–24]. A number of third-generation calcium indicators have since been developed with a wide range of excitation spectra and calcium affinities, including the currently popular dyes Oregon-Green BAPTA (OGB) and fluo-4 [25], whose fluorescence increases in line with neuronal calcium elevations upon two-photon excitation. This property, particularly advantageous in noisy preparations such as those experienced in vivo, has contributed to their rise in popularity to become the dyes of choice today [e.g. 26–28]. However, greater SNR has been achieved by two new synthetic dyes, Cal-520 and Cal-590 [29, 30], which are able to reliably detect single action potentials in both brain slice and whole brain preparations, whilst remaining relatively simple to load. Cal-520 is able to detect fourfold the number of active neurons in barrel cortex in vivo when compared to OGB (see Fig. 1, and Table 1), and can reliably detect individual action potentials in 10 Hz spike trains [29], therefore improving the achievable spatiotemporal fidelity of neuronal network interrogation. Cal-590 is a red-shifted synthetic dye, enabling previously inaccessible neuronal calcium signals to be assessed in all six layers of cortical depth as scattering of photons at longer wavelengths is reduced. At up to 900 *μ*m beneath the pial surface, imaging can be performed with Cal-590 while maintaining good signal-to-noise ratio and preserving spatiotemporal fidelity [30].

It is important to note that whilst the ability to bolus load synthetic dyes represents a distinct advantage for ease of use, these third-generation indicators are available in both membrane-permeable and membrane-impermeable versions, enabling their use in a variety of different experimental paradigms. Perfusion of membrane-impermeable dyes through an intracellular recording pipette enables single action potentials, and individual synaptic inputs, to be resolved in both in vitro and in vivo preparations [29, 30, 43–48]. Bolus loading of membrane-permeable dyes enables hundreds of neurons to be examined simultaneously [18,29, 30,49]. There are however several costs to this. The first is that background fluorescence is relatively high (and labelling is not restricted to neurons), meaning that it can be difficult to resolve individual neuronal processes. Additionally, dye loading lasts only a matter of hours, meaning that it is not possible to use this approach for chronic neurophysiological investigations.

**Fig. 1.**
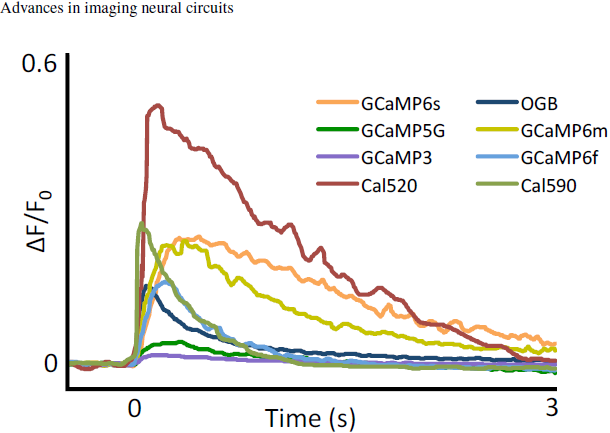
Comparison of calcium sensitive indicators. Responses averaged across multiple neurons and wells for GCaMP3, 5G, 6f, 6m, 6s, OGB, Cal-520 and Cal-590; fluorescence changes in response to 1 action potential. Figure adapted from [29–31]

### 2.2 *Calcium-sensitive genetically encoded proteins*

The invention of protein-based genetically encoded calcium indicators (GECIs) [50] represents an important milestone in the field of biosensors. One main advantage of GECIs over synthetic dyes is that these proteins can be targeted to specific cell types [51–54], enabling non-invasive imaging of neuronal networks [55–58] over chronic timescales [59].

Among the GECIs, those based on the GCaMP framework, consisting of circularly pennutated GFP fused to calmodulin and a Ca2+/calmodulin-binding myosin light chain kinase fragment, are the best studied [60, 61]. The early types of GCaMP proteins had limited application due to their slow response kinetics and low signal-to-noise ratios in comparison to OGB, meaning that neuroscientists had to compromise between highly sensitive probes delivered via invasive injection methods, or GECIs with poor sensitivity delivered by non-invasive genetic methods. Fortunately, following multiple rounds of structure-guided design [53, 62, 63], development of the GCaMP6 series represents the first GECIs with higher sensitivity than the currently most commonly used synthetic dye, OGB [31]. In addition, there have been recent efforts focused upon the development of red fluorescent GECIs (RCaMPs) [62, 63]. RCaMPs, which enable imaging to be performed in deeper tissue [64], provide scope to achieve dual-color fluorescence microscopy in combination with green GECIs. R-CaMP2 holds particular promise for in vivo applications, possessing both strong single action potential signals, and fast kinetics [42].

**Table 1.** Comparison of the properties of some frequently used calcium indicators. Higher dissociation constant *K_d_* implies better temporal fidelity of the dye, at the expense of accuracy in readout of baseline calcium levels. Shorter rise time is desirable for accurate detection of action potential onset timing. Shorter decay time may allow single action potentials to be followed from higher firing rate spike trains, but longer decay times lead to better detectability of single action potentials.

| Name | $K_d$ (nM) | Rise Time | Decay Time Constant | References |
| --- | --- | --- | --- | --- |
| **Synthetic Calcium Dyes** |  |  |  |  |
| Aequorin | $10^4$ | | | [32] |
| Arsenazo III | $10^7$ | | | [33] |
| Quin-2 | 89 |  |  | [34] |
| Fura-2 | 140 | <20 ms | <40 ms | [35, 36] |
| Indo-1 | 230 | < 100 ms | < 250 ms | [37, 38] |
| Fluo-3 | 390 | < 10 sec | < 10 sec | [39] |
| Oregon Green BAPTA-1 | 170 | <100 ms | 1-3 sec | [29, 40] |
| Fluo-4 | 345 | < 80 ms | < 250 ms | [40] |
| Cal-520 | 320 | <70 ms | < 800 ms | [29] |
| Cal-590 | 561 | <2 ms | < 30 ms | [30] |
| **Calcium Sensitive Genetically Encoded Proteins** |  |  |  |  |
| GCaMP3 | 365 | < 100 ms | < 1 sec | [31, 41] |
| GCaMP5G | 460 | < 200 ms | < 1 sec | [31, 41] |
| GCaMP6f | 296 | < 200 ms | < 1 sec | [31] |
| GCaMP6s | 152 | < 300 ms | > 3 sec | [31] |
| R-CaMP2 | 69 | < 60 ms | < 600 ms | [42] |

GECIs have several advantages over synthetic dyes in that they can be targeted to specific cell types, and their chronic expression enables the same neurons to be imaged during multiple sessions over a long time scale. However, synthetic dyes still have advantages in acute experiments due to ease of labelling, a relatively linear calcium response, and high sensitivity.

### 2.3 *Membrane potential sensors*

Although calcium indicators can be an effective means of resolving sparse (low firing rate) spike trains, their slow kinetics (and the slow kinetics of the underlying calcium changes in cells) make them non-ideal for observing spike trains from neurons with higher firing rates. In addition, calcium signals relating to the action potential are confounded by complex internal calcium dynamics. A reasonable alternative is therefore the direct imaging of membrane voltage fluctuations. Organic voltage sensitive dyes provide one means of doing this, and have proved useful in some cases for mesoscopic scale imaging of neural activity [e.g. 65]. Given that the action potential has a duration on the order of 1 ms, however, most of the signal observed through mesoscopic membrane potential imaging is subthreshold. Organic dye membrane potential reporters have relatively poor signal to noise ratios, and have not been found to be capable of resolving single action potentials in-vivo [66]. In slice, however, Single-cell styryl dye loading through whole-cell patch pipettes has enabled single-trial imaging of action potentials in CNS dendrites and spines[67], axons [68, 69], collaterals and presynaptic boutons [70], as well as post-synaptic potentials in dendritic spines [71]. Phototoxicity effects limit the applicability of this technique. Recently, Roger Tsien’s group recently developed a fast, high-sensitivity voltage sensor based on photo-induced electron transfer [72]. In principle, the enhanced-sensitivity probes should require lower light intensities to achieve high S/N recordings thus reducing photodamage. However, their utility has not yet been demonstrated in intact slices.

Genetically encodable voltage indicators (GEVIs), however, at least partially resolve these issues, allowing genetically targeted neurons to be monitored over chronic timescales [73]. There are two main classes of GEVIs [reviewed in 74], voltage sensitive fluorescent proteins (VSFPs), which make use of a voltage sensing domain and microbial opsin based GEVIs. There have been numerous developments in both families in recent years. Voltage sensing domain based sensors have been improved with the development of a number of variants of the VSFP family (VSFP2s, VSFP3s, VSFP-Butterflies) [74], ArcLight, and recently the highly sensitive ASAP1 [75]. At the same time, work progressed on voltage sensors based on the rhodopsin Arch, with modifications based on mutation and designed manipulations yielding sensors with improved kinetics and sensitivity, although still low quantum yields [76]. Most recently, a voltage-sensitive fluorescence resonance energy transfer (FRET) based sensor family, Ace-mNeon, was developed which combines both the fast kinetics of a rhodopsin voltage sensitive domain with a bright fluorophore. The result is a membrane potential sensor (Ace2N-mNeon) which appears to be capable of single trial detection of individual actions *in vivo* [77]. Antic et al. recently published a detailed review of GEVI developments [78].

## 3 Galvanometric scanning approaches

The standard scanning approach used in two photon microscopy is to use a moving magnet galvanometer to control the position of a mirror in the optical path. We refer to this as a Galvanometric Scanner (GS). Driving two GSs in specific motions allows the user to direct the laser beam through different trajectories over a neural circuit loaded with activity-sensitive fluorophores (see Section 2). Trajectories can be optimised either for resolving the structure of the neural circuit, or for obtaining fine temporal resolution acquisition of functional signals. Galvanometric scanning is inherently a sequential technique (but see beamlet discussion in Section 5), and thus is ultimately unlikely to scale well to mesoscopic analysis of neural circuits. However, galvanometric scanning is a mature technology, and in the short to medium term, may provide the most practical way of scanning populations of up to approximately one thousand neurons. In this Section, we describe different galvanometric scanning approaches for two photon microscopy.

**Fig. 2.**
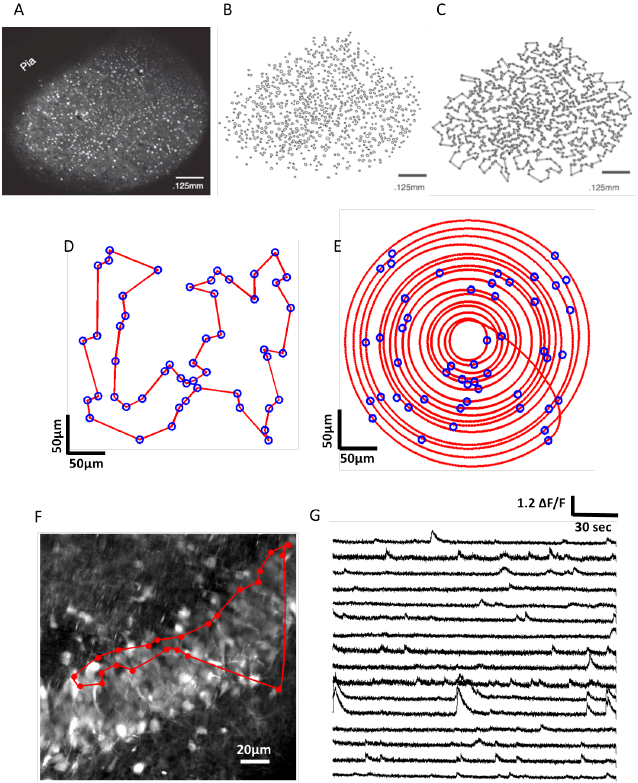
Galvanometric scanning approaches. A)-C) Steps involved in the HOPS algorithm: extraction of cell body locations, followed by approximate solution of the shortest path (Travelling Salesman problem) (A,B and C come from the publication: Heuristically optimal path scanning for high-speed multiphoton circuit imaging; Sadovsky AJ, Kruskal PB, Kimmel JM, Otermeyer J, Neubauer FB, MacLean JN; J. Neurophysiology Sept. 2011 106(3):1591-8). D) Travelling salesman solution for an ensemble of 50 cells in a 300 × 300 *μ*m region of simulated tissue. E) Adaptive spiral scanning solution for the same population of cells. F) High resolution (frame-scanned) image of the granular layer of the dentate gyrus: The red circles represent the 23 detected spontaneous active neurons during raster scanning and the red trajectory represent the traveling salesman path followed by the laser beam. G) Calcium traces sampled at 420 Hz with a Traveling Salesman trajectory passing through 16 selected of 23 active neurons.

### 3.1 *Galvanometric scanners*

Most modern galvanometric scanners share a common architecture: a moving magnet torque motor, a mirror and an optical position transducer. For the purpose of microscopy, two types of GS are used: standard and resonant GS (RGS). Regular GSs may be driven by any kind of waveform, subject to with frequency and amplitude limitations, whereas RGS can only be driven by a sine wave at their resonant frequency. Several optics manufacturers, including Cambridge Technology and Thorlabs, sell both types of GS; standard models for two-photon microscopy in use in our laboratory are the 6215H (standard) and CRS8kHz (resonant) from Cambridge Technology.

### 3.2 *Raster scanning*

With Regular GSs, the most common scanning method is raster scanning (see Fig. 2). This consists of driving the laser beam in tight parallel lines over a rectangular area, to build up an image of the labelled tissue. For standard galvos, either sawtooth or triangular waveforms are typically used, with the former being simpler for image reconstruction, and the latter enabling twice the repetition rate for the same power consumption. For both driving waveforms, the critical point is the point where direction is reversed, as it contains higher frequencies and can induce heating in the system. As the extreme parts of the field of view (FOV) are not crucial for image acquisition, another approach that can be taken is to use a segment of a sine wave that matches the slope of the linear portion of the signal, and use a Fourier decomposition of those driving waveforms. In [79] it was demonstrated that the capabilities of raster scanning can be enhanced by a Fourier decomposition approach where the coefficients are chosen carefully to minimize the deviation from linearity during the acquisition period. They report a sampling rate of around 100*Hz* covering a very small scan area of 8.25 × 4.75 *μ*m^2^.

Resonant galvos can also be used for raster scanning, when coupled with a regular GS. RGS, typically with resonance frequencies from 4 to 12 kHz, are assigned to the fast scan axis, with a standard GS on the slow axis [80]. In this configuration, a typically achievable framerate (in our hands) for an image of 300 × 300 *μ*m^2^ at a resolution of 512 × 512 pixels, a resonant frequency of 8 kHz, and a pixel dwell time of 4 *μ*s is approximately 30 Hz; the framerate can be increased in multiples by restricting the field of view. While resonant scanning allows framerates to be increased, in some cases to impressive values, it suffers from the same issue as other raster scanning approaches: much of the photon collection time is wasted scanning structures of little or no interest.

### 3.3 *Scanning algorithms for multiphoton microscopy*

The development of novel algorithms for scanning is motivated by bandwidth constraints: two photon microscopy is essentially a “point” measurement technique, and as most points within an image are not of interest for reconstructing a neural circuit, bandwidth can most effectively be employed by constructing trajectories focused on points of interest. This is analogous to the saccadic eye movements with which the human visual system scans a visual scene.

Novel scanning strategies offer an accessible approach to increasing the temporal resolution of two photon microscopy. They can be implemented on most commercial or custom two photon microscope with minimal hardware changes; adapted control software is usually all that is required.

One early development of such a scanning strategy was achieved by Lillis et al.
[81], who were able to record calcium traces from points across an entire rat hippocampus at scan rates of 100 Hz. After performing a Raster scan, the user needed to select multiple segments of interest (SOI) in the image. These SOIs were then linked to create a closed scanning path. Moreover, their algorithm controlled the scanning speed such that the acquisition was performed done at constant velocity within the SOIs and with maximum acceleration and deceleration between the SOIs. This scanning strategy demonstrated substantial advantages in comparison to raster scanning particularly in the situation of sparsely distributed neurons across a large region.

### 3.4 *Heuristically optimal path scanning (HOPS)*

Sadovsky et al. [82] developed a scanning algorithm (Heuristically Optimal Path Scanning, or HOPS) based on the Lin-Kernighan Heuristic Traveling Salesman Algorithm. After performing a regular raster frame scan on a mouse somatosensory cortex slice loaded with the inorganic dye Fura-2 AM, to obtain a high resolution image of the labelled tissue, they detected cell body centroids using custom-developed software. They then used the LKH algorithm to find a nearly optimal order for scanning the cells in order to minimise the path distance (see Fig 2C). HOPS additionally allows the dwell time over a cell, before moving on to the next cell, to be set. This enables a balance to be achieved between maximising sampling rate and SNR, depending upon specific experimental needs. With this scanning strategy, they achieved sampling rates as high as 150 Hz for 50 cells and around 8.5 Hz for 1000 cells.

This approach has been used successfully to study the spatiotemporal dynamics and functional connectivity of cortical circuits *in vitro* [e.g. 83]. After frame scanning the granular layer of the dentate gyrus in mouse hippocampal slices loaded with the synthetic calcium dye Cal-520, we used the Traveling salesman algorithm and show calcium traces sampled at 420*Hz* from sixteen of the twenty three spontaneous detected active neurons (see Fig 2 F and G). However, it should be noted that such an approach does not take into account the kinematic properties of the galvanometer: a truly optimal approach would incorporate the momentum of the mirrors into the choice of trajectory.

### 3.5 *Adaptive spiral scanning*

The main drawback of scanning strategies that do not take into account the inertia of the GS is that their trajectories contain high frequency components that will be filtered when the GSs are driven at high velocity. This limits the sampling rate that can be achieved to precisely follow every sharp turn of the path. Hence, an alternative approach is to recognise that the fastest trajectory that can be achieved by a pair of mirrors is elliptical (or circular, in the case of identical mirrors). By changing the radius of the elliptical trajectory, spiral patterns can be achieved. This is recognized in the approach taken by the Adaptive Spiral Scanning (SSA) algorithm [84], which finds a set of spirals traversing the cell bodies of interest to maximise sampling rate. This theoretical model, suggests that sampling rates of 475 Hz could be achieved for 50 cells, and 105 Hz for 1000 cells, if the galvo is driven in open loop mode at 10 kHz, well within its dynamic range [84]. However, galvanometric scanners are typically controlled in closed loop mode with a PID controller, with an allowable driving frequency of approximately 1 kHz, depending upon amplitude. Under these conditions, performance is reduced; in real world tests using a driving frequency of 1 kHz with 65 targets (fluorescent beads), the maximum sampling rate that could be achieved with the SSA was 103 Hz. For 152 targets, the maximum sampling rate achievable was 63 Hz. In any case, it is apparent that with customised galvanometric control hardware (i.e. either operating in open loop or in closed loop but with a PID controller optimised for the dynamics of spiral trajectories), it is possible to create desired trajectories at sampling rates exceeding those achievable by the HOPS algorithm. Adaptive spiral scanning can also be naturally extended to three-dimensional scanning strategies through the incorporation of a fast z-axis scanner such as an Electrically Tunable Lens [85].

## 4 Acousto-optic scanning

The GS’s inertia inherently limits the speed at which they can change position. Inertialess systems, such as Acousto-optic deflectors (AODs), can offer much higher switching speeds [86].

AODs diffract light from periodic refractive index variations in crystals caused by acoustic waves. The ultrasonic acoustic frequency governs the light diffraction angle. Orthogonal AOD pairs allow random access of any point in the *x*-*y* FOV in short times (~ 10 *μ*s, Grewe and Helmchen [87]).

Fig. 3 shows a typical AOD configuration. If the beam is incident on the AOD at the Bragg angle then only the efficiency of diffraction to the first order is maximised [88]. The first order deflection angle, *θ* is then given by [89]:

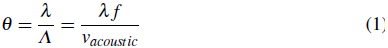

Where λ is the optical wavelength, *Λ* the acoustic wavelength, *f* the acoustic frequency and *v_acoustic_* the acoustic wave speed. The angular AOD scan range, Δ*θ_scan_* is therefore:

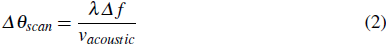

Where *Δf* is the AOD acoustic bandwidth. A common parameter for objective scanning technology comparison is the number of resolvable spots, *N*, defined as

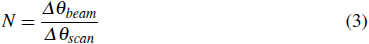

where *Δθ_beam_* is the excitation beam divergence. This can be interpreted as the deflector’s resolution as it is the number of independent points that can be sampled.

Acoustic waves commonly propagate in crystals in one of two modes: fast longitudinal compression waves and slow shear waves [23]. AODs are most used in shear mode as it supports an increased acoustic bandwidth that allows increased spatial resolution [89]. As the deflection switching time is dependent upon the time it takes the acoustic wave to fill the crystal, using this mode does decrease the temporal resolution.

Spatial and temporal dispersion is a major consideration in any multiphoton AOD scanning implementation [94]. 2P imaging requires ultrashort pulsed lasers for excitation. Short pulses necessarily have a large spectral range due to the constraint on the time-bandwidth product. Different pulse spectral components separate in space and time as they diffract and propagate through an AOD. Spatial dispersion distorts the system’s point spread function (PSF), increasing *Δθ_beam_* and reducing the resolution by at least 3x for standard imaging conditions [89]. Temporal dispersion limits the pulse peak power, leading to a drop in collected fluorescence and degraded SNR. Additional diffractive optical elements can mitigate these effects by introducing opposing dispersion into the beam path to compensate. This can be achieved using a single prism [95] or by multiple element systems [91]. AODs, along with some dispersion compensation components, introduce large power losses into the imaging set-up [96]. This can cause the maximum imaging depth to be power limited [91].

AODs provide a small maximum deflection angle, *Δθ_scan_* typically 0.01-0.05 rad compared to 0.5-1 rad for a galvanometer scanner [97]. This limits the image resolution to the AOD pointing accuracy if a large FOV is required [87].

Acousto optic lenses can also be constructed using chirped acoustic waves to drive AOD pairs [90], as depicted in Fig. 3B. These can simultaneously scan in *x, y* and *z* by deflecting the laser beam whilst causing it to converge or diverge slightly.

**Fig. 3.**
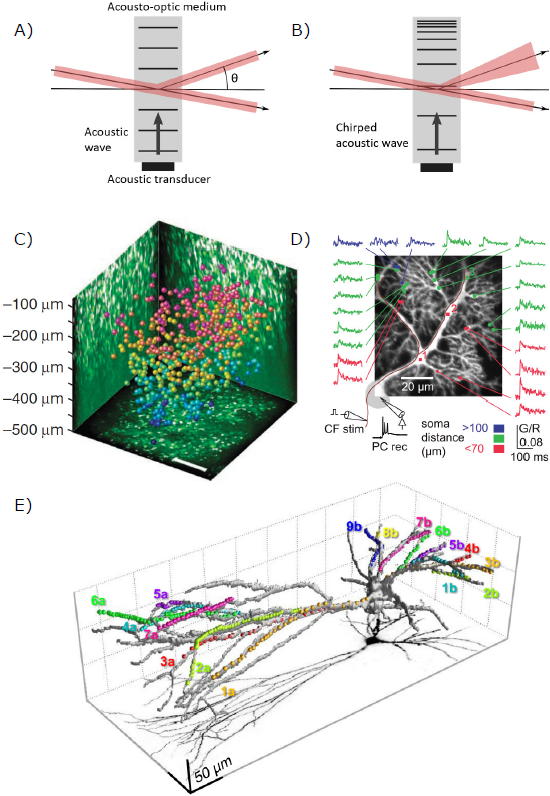
Acousto-optic scanning. A & B) Two methods of using AOD scanners. A) Unchirped acoustic waves diffract light allowing 2D scanning. B) Chirped acoustic waves diffract light whilst causing it to diverge or converge, allowing scanning in 3D. Adapted from Reddy and Saggau [90]. C) Katona et al. [91] were able to scan 532 neurons in a 400×400×500 *μ*m FOV that at ~ 56Hz. Reprinted by permission from Macmillan Publishers Ltd: Nature Methods. ref. [91], copyright 2012 D) Purkinje cell dendritic tree showing quasisimultaneuos calcium imaging of response to climbing fibre stimulation. From ref. [92] licensed under CC BY 3.0. E) 3D Scanning of a fast spiking PV interneuron [93]. Dots show locations of sampling for Calcium imaging.Reprinted from Neuron, 82(4), Balzs Chiovini, Gergely F. Turi, Gergely Katona, Attila Kaszs, Dnes Plfi, Pl Mak, Gergely Szalay, Mtys Forin Szab, Gbor Szab, Zoltn Szadai, Szabolcs Kli, Balzs Rzsa, Dendritic Spikes Induce Ripples in Parvalbumin Interneurons during Hippocampal Sharp Waves, 908-924, Copyright 2014, with permission from Elsevier.

This causes the beam to focus above or below the objective nominal focal plane, although at a cost to the maximum lateral deflection angle, allowing for random access to a diamond shaped 3D FOV [98]. This scheme also inherently compensates for spatial dispersion, reducing the compensation required in the beam path.

A 3D AOD scanning implementation combining chirped AODs for axial and unchirped for lateral focussing has achieved a large 700*μ*m × 700*μ*m × 1,400*μ*m maximal square FOV by using detailed optical modelling and extensive dispersion compensation [91].

Addressing the technical challenges generally requires a microscope specifically designed for AOD scanning. Unlike with novel galvanometric scanning strategies, commercial or custom microscopes would require major adaptations in order to fully exploit the advantages of AODs.

High speed AODs have enabled multi-cell imaging over large areas [99] as well as quasi simultaneous monitoring of dendritic spine synaptic input [100]. Dendritic imaging of fast spiking parvalbumin interneurons allowed sharp wave ripple to be imaged [93]. AOD scanning has also been extensively used for monitoring Ca^2+^ transients from randomly accessed regions of Purkinje cell dendritic trees [88, 92, 101].

## 5 Parallel scanning and scanless methods

The preceding sections detail efforts to accelerate functional fluorescence image acquisition through AOD, galvanometric or resonant scanning of a single diffraction limited spot. Here we discuss complementary technologies to increase speed, or number of targets sampled, by reshaping the focal volume to interrogate multiple locations in parallel.

### 5.1 *Multiple beamlets*

Efforts to increase functional fluorescence acquisition speed have evolved optical systems that divide the laser beam into several beamlets with spinning disk [102] and square xy-scanned [103] beamlet arrays, as well as etalons [104] and cascaded beam splitters [105, 106]. Importantly, acquisition rates increase in proportion to the number of beamlets. Moreover, due to nonlinear photodamage [107], the power density at focal volume for single beam scanning is typically limited to a small fraction of the total power available. Multifocal schemes enable increased fluorescence photon rates while decreasing the photodamage probability as any single focus. Acquisition speed can thus increase without sacrificing signal-to-noise ratio while conserving optical sectioning similar to single-beam scanning.

The multiple simultaneous foci necessitate fluorescence detection with a spatially resolved detector, often a charge coupled device (CCD) camera. This requirement imposes a critical limitation for neuroscience applications: the imaged fluorescence cannot be unambiguously assigned to a focus deep in scattering brain tissue. Innovations to reduce cross-talk between adjacent foci include temporal pulse multiplexing [108], confocal filtering [109], and mapping of foci onto large detector multianode PMTs [110]. Nonetheless, maximum imaging depths in scattering mammalian brain tissue to date range from 50 to 100 microns, inferior to the many hundreds of microns depth penetration achieved with single beam scanning.

### 5.2 *Multi-beam multiple fields of view*

Two-photon microscopes typically feature single lateral fields of view (FOVs) spanning 500 ± 200 *μ*m. Although useful for single-cell interrogation at the microcircuit scale, this FOV is too small to observe functions spanning multiple brain areas (e.g., sensory-motor integration, decision making).

Lecoq et al. developed a dual-axis microscope enabling simultaneous, independent imaging of two spatially separated regions [111]. They positioned gradient-index (GRIN) microendoscope doublets (1 mm diameter, 0.35 relay NA) over the two regions of interest, one coupled to a miniature mirror folding the optical path by 90° with respect to the vertical axis. Two standard long working distance objectives couple the GRIN endoscopes to two separate fluorescence detection and excitation pathways supplied by a single Ti:Sapphire laser split prior to the galvo scanners. The system can image two 700 *μ*m FOVs, either adjacent or separated by up to several millimeters. The point spread function (1 *μ*m lateral, 9 *μ*m axial), limited by the low effective NA (0.35), nonetheless enabled high signal-to-noise single-cell resolution calcium fluorescence imaging with which they simultaneously monitored primary and latermedial visual cortices.

The quest to marry circuit activity monitoring across micro-and meso-scales recently spurred development of mega-FOV two-photon microscopes. Importantly, two groups achieved 5-to 10-fold increased lateral FOVs primarily through careful astigmatism correction for large scan angles [112, 113]. In order to increase frame rates over two simultaneously acquired FOVs, Stirman et al. [113] split the beam, sending one component through a delay arm beam before guiding both beams through independent, galvo-based x/y scanners. The femtosecond pulses from the delay and short arms interdigitate in time and can thus be temporally demultiplexed at the shared detector, in order to assign collected photons to the correct field of view. This scheme overcomes one critical limitation to Lecoq et al. [111]’s dual-region microscopy, in that the positions of the two ROIs are flexible across the mega-FOV, not fixed beneath surgically implanted endoscopes.

In addition to lateral FOV separation, the temporal pulse multiplexing method described above has accomplished multiple simultaneous axial plane imaging [114]. Here, the laser beam splits to introduce differing divergences and delays such that, when recombined into the scan path, focus into separate axial planes, with temporally interdigitated pulses. Cheng et al. [46] integrated this system with a resonant scanner to achieve framerates of up to 250 Hz across 400 × 400 micron FOVs, simultaneously in four planes spanning mouse cortical layers 2/3.

To conclude, in contrast with the multiple beamlet techniques discussed in the previous section, the signal arising in each location can be unambiguously assigned to a specific excitation beamlet, either by spatial separation[111] or temporal multiplexing[46, 113]. Scattered photons incident on a point detector thus contribute to useful signal rather than background contamination, enabling high contrast images at depth in scattering tissue.

### 5.3 *Light sheets*

Light sheet microscopy achieves optical sectioning by squeezing fluorescence excitation into a plane [115, 116] or line [117] orthogonal to the imaging axis. The sample translates with respect to this plane in order to build up a three dimensional image. Importantly, light-sheet enables whole line or plane illumination, rendering image acquisition “scanless” in one or two spatial dimensions. Compared to point raster scanning, simultaneous line or plane imaging reduces the frame period by a factor *n* or *n*^2^ respectively (for a *n* × *n* sized image), without sacrificing fluorescence photon flux. Notably, Ahrens et al. [118] recently capitalized on this speed advantage to image spike-correlated calcium transients simultaneously from 80% of the neurons in a larval zebrafish brain with single-cell resolution (Fig. 4).

Light sheets are typically generated by focusing a laser beam in one direction with a cylindrical lens [119, 120]. The finite distance over which a Gaussian beam can focus to a thin waist forces a compromise between the sheet thickness and lateral FOV size. Alternatively, a circular beam scans in one direction, sweeping out an excitation plane (Fig. 4A). Investigators recently increased FOV size by scanning Airy beams [121] or Bessel beams [122], that unlike Gaussian profiles, do not vary in intensity along the propagation axis.

Palero et al. [123] implemented the first two-photon light-sheet microscope with cylindrical lenses that focused the laser beam into a sheet. Two-photon excitation improves resolution and depth penetration over one-photon light sheets, owing to decreased scattering at long wavelengths and the signal’s quadratic dependence on the excitation intensity. However, this same nonlinearity drastically decreases fluorescence excitation levels with low power densities due to distribution of the laser beam into a sheet. Keller et al. [117] improved on this by laterally scanning a spherically focused beam, reporting 100-fold increase in fluorescence.

The orthogonal orientation of illumination and imaging axes would seem to limit light-sheet methods to small, transparent samples such as zebrafish and C. *elegans*. However, Dunsby [124] developed a system that illuminates and images an oblique light sheet through a single high NA objective, paving the way for a swept version devised to acquire rapid volumetric images in preparations such as neocortex that preclude side on illumination geometries. Impressively, Bouchard et al. [125] demonstrated calcium transient imaging in awake behaving mouse dendrites at rates exceeding 20 volumes per second. Although developed for one photon excitation, this configuration is amenable to two photon implementations able to reduce scatter and increase penetration depth.

**Fig. 4.**
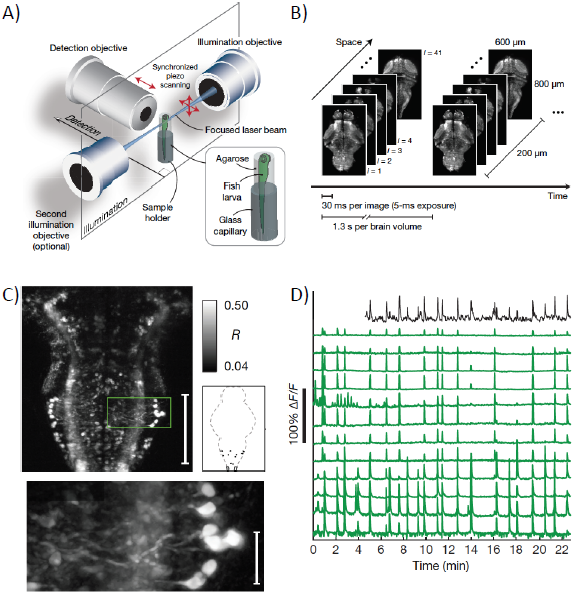
Whole brain, single-cell resolution calcium transients captured with light-sheet microscopy (reproduced with permission from Ahrens et al. [118]). (A) Two spherically focused beams rapidly swept out a four-micron-thick plane orthogonal to the imaging axis. The beam and objective step together along the imaging axis to build up three-dimensional volume image at 0.8 Hz (B). (C) Rapid light-sheet imaging of GCaMP5G calcium transients revealed a specific hindbrain neuron population (D, green traces) traces correlated with spinal cord neuropil activity (black trace).

Oron et al. recently pioneered an alternative technique, temporal focusing, that enables two-photon line-or sheet-generation on-axis. Briefly, they manipulate the temporal profile of femtosecond pulse such that it achieves minimum duration and maximum peak power only at the plane of focus, stretching out before and beyond. Two-photon fluorescence is thus excited with high probability only at the “temporal focal plane,” in the range of 1-10 microns thick. Temporal focusing can be combined with other scanning methods for calcium imaging [126], however its real advantages may lie in widefield application. Oron and colleagues demonstrated depth resolved images built up from temporally focused lines [127] and sheets [128], also known as ‘’temporally focused widefield two-photon.” Since then, others have combined temporally focused sheets with piezo-[129] and ETL-based [130] axial scanning for rapid volumetric imaging of calcium transients in zebrafish and C. *elegans*. Since temporally-focused light sheets and lines generate in planes orthogonal to the imaging axis, their implementation for imaging mammalian brains *in vivo* may prove more simple than Bouchard et al. [125]’s obliquely generated light sheets, although this remains to be demonstrated.

### 5.4 *Holographic excitation*

The methods discussed above assume that all locations in the line-or sheet-swept area bear equal information content. In cases where locations of cells or structures or interest are known, one can gain further by targeting excitation light exclusively to these regions. Computer-generated holography (CGH) has emerged as a convenient method for distributing light across targets with high spatial and temporal flexibility.

CGH shapes light at the sample plane by manipulating the phase of an expanded laser beam at the microscope objective back aperture. Phase modulation is accomplished with reflective spatial light modulators (SLMs) arrays, typically liquid crystal on silicon (LCOS), occupying a plane conjugate to the objective back focal plane. Investigators first image target cells or structures, then design a mask or series of masks over which light should be patterned. The corresponding phase patterns addressed to the SLM are calculated using algorithms based on gratings and lenses [131] or iterative Fourier transformation [132, 133], featuring differing degrees of simplicity, flexibility and uniformity. While SLM refresh rates currently limit the rate at which patterns can change to 10s to 100s of Hz, new developments such as high-tilt ferroelectric liquid crystal analog modulation may soon enable kilohertz switching rates [134] similar to commercially available binary (0, *π*) devices. Holographic shape axial extent scales linearly with lateral area resulting in loss of resolution in this axis for large targets. This can been mitigated by combining two-photon CGH with temporal focusing to decouple axial confinement from lateral extent [135]. Two-photon CGH-TF also improves depth penetration and renders shapes more robust to scattering [136].

In addition to CGH’s growing popularity for controlling neural circuits [reviewed in 137], and its potential for studying single cell function [138], recent studies demonstrate its utility for scanless, multi-target fluorescence imaging. Nikolenko et al. [139] targeted multiple cortical neurons *in vitro* with two-photon CGH-generated spots to image calcium transients in 20-50 cells simultaneously at 15-60 Hz. Ducros et al. [140] captured cortical calcium fluorescence transients *in vivo* with multiple CGH spots simultaneously generated in multiple planes at depths up to 300 microns. In order to discriminate between signals arising from the multiple spots with a single PMT, they added a temporal encoding scheme, modulating each spot in time with a digital micromirror device (DMD). Foust et al. [141] recently adapted CGH-based imaging to achieve high signal-to-noise, fast voltage imaging in cortical slices. In contrast with the spot illumination utilized by Nikolenko and Ducros, they patterned light over extended regions of axons and dendrites, enabling improved spatial specificity compared to wide-field illumination. Importantly, extended shapes increase spatial specificity while limiting compromises in speed or signal-to-noise ratio compared to diffraction-limited point scanning.

Despite notable gains in speed, spatial specificity and photodamage reduction achievable with light sheet and holographic methods, the necessity to image emitted fluorescence, as with multifocal multiphoton microscopy, severely limits the depth at which high contrast images can be obtained in scattering tissue (demonstrated up to 134 microns [125]). Tissue-specific factors including scattering length (e.g., longer for younger, less process-and myelin-dense regions) and the spatial sparseness of the fluorescent neurons critically determine this depth limit. Red shifted fluorophores [30, 142], structured illumination and post-processing strategies [143, 144] may soon improve on current depth limits. In the future, light-sheet and holographic techniques can be combined with “micro-optics” such as glass plugs [145], gradient-index lenses[146], andmicroprisms[147] to access structures beyond limits imposed by scattering. Lightsheet and CGH techniques require extensive adaptation of traditional epifluorescence microscopes as an additional dichroic must be used to introduce these distinct beam paths.

## 6 Implications for the development of data analysis tools

### 6.1 *Pre-processing of two photon imaging data*

The rapid and substantial advances in technologies for experimental neuroscience described above require equally substantial developments in data analysis tools. However, both signal processing and data analysis tools for multiphoton imaging are in a relatively rudimentary state in comparison to those available for single unit and multi-electrode array electrophysiology, which has had decades of development.

A first basic step in analysis of two photon imaging data (particularly when twodimensional scanning approaches such as raster scanning are used) is the segmentation of regions of interest (ROIs). This is necessary to pool photons from locations producing the same signal, in order to obtain sufficient SNR for further analysis. Until recently, ROI segmentation has in many cases been performed manually. However, manual segmentation does not scale well, and with some instances in which it has been possible to record from very large numbers of neurons [118], the development of robust automated approaches is essential. The difficulty of this varies substantially according to the fluorophore and load/labelling technique used. It is relatively straightforward to obtain somatic ROIs from cortical tissue loaded with AM-ester dyes such as OGB-1 AM [148], however dyes with lower baseline fluorescence (such as Cal-520 AM) may benefit from improved ROI segmentation procedures. In addition, optimal ROI segmentation may result in higher SNR due to rejection of optical cross-talk due to signals from nearby structures. In some brain areas, such as the cerebellum, the task is more difficult, due to specific morphological properties of cells. A number of methods have therefore been proposed to improve ROI segmentation, including spatial independent component analysis [28, 149, 150] and pixel-wise cross-correlation followed by anisotropic or morphological filtering [28,151]. A more advanced approach may be to simultaneously find the spike trains and the regions of interest that generate them, using approaches such as structured matrix factorization [152, 153]. This can provide one way to demix pixels from overlapping ROIs (which may occur, for instance, when processes from two neurons pass within an optical point spread function of each other). Good solutions for the ROI demixing problem may improve the SNR of somatic ROIs also, by removing neuropil contamination without “wasting” potential signal power from boundary pixels.

Having obtained a time series for each detected neuron, a common second stage of processing is to perform event detection-turning a time series comprised of calcium transients into a spike train. This places the data on the same terms (although not quite the same temporal resolution) as extracellular electrophysiological data. Given the kinetics of the calcium signal and the relevant fluorophores, this process is much more accurate at low firing rates. Where firing rates are higher, it may be more useful to work with continuous signals-either the original calcium signals [148], or with signals to which a deconvolution filter has been applied [154]. In this latter approach, the aim is to invert the linear filter operator between the calcium signal and the ground truth spike train, recorded electrophysiologically for a sample neuron. This inverse filter can then be applied to all calcium time series, producing an estimate of instantaneous firing rate in each cell. This is necessarily approximate, as it neglects inter-cell variability-the calcium dynamics of any given cell may differ from the cell used to find the filter, and thus the signal for some cells may be distorted with respect to the instantaneous firing rate interpretation. A simple alternative is to apply a rectified smoothing derivative filter (which the deconvolution (inverse) filters strongly resemble in any case).

For many applications, ranging from receptive field mapping [151] to crosscorrelation analysis [28], continuous time series calcium signals may be appropriate. However, the action potential is the means by which information is transmitted from one neuron to another, and neuronal calcium dynamics reflect other processes than just those due to action potential related calcium influx. The detection of calcium transients is thus an important problem. Early approaches to this problem made use of template matching, with either fixed templates [155] or templates derived from data [28]. A variant of this is the “peeling” algorithm: successive template matching, followed by subtraction of the matched template from the data [156]. This is one way to deal with the problem of having multiple overlapping calcium transients (provided that they add linearly-we have found this to be true in some cases but not in others). However, it is not clear that the high computational demands of this approach pay off in performance terms.

We recently introduced a novel approach to calcium transient detection, based on Finite Rate of Innovation (FRI) theory. FRI theory considers signals that have a finite number of degrees of freedom per unit time [157], and provides methods for sampling and reconstructing such signals with a kernel appropriate to the characteristics and information content of the signals. In this case, we know that the shape of calcium transients tends to follow a single exponential waveform; thus the problem reduces to reconstructing a stream of decaying exponentials [158]. This results in an algorithm which is fast, non-iterative, parallelizable, and highly accurate. The approach is model-based, but with the strength that it has polynomial complexity, whereas true template matching approaches require a combinatorial search in order to achieve the same performance.

To validate our approach, we performed two photon targeted electrophysiological recordings from the dendrites or somas of cerebellar Purkinje cells, while imaging calcium signals in the vicinity [Fig. 5A-B; 28, 158]. This was used to generate surrogate data of controlled SNR (Fig. 5C). In subsequent work, the FRI calcium transient detection algorithm was modified to specifically take account of GCaMP6 kinetics, as well as improving the way it dealt with multiple overlapping events [159, 160], further improving performance.

One approach to the calcium transient detection problem is to use a deconvolution procedure (as described above), followed by a thresholding process to infer a resulting spike train [161]. Despite this being perhaps the most common method currently in use, we did not find it to perform particularly well (Fig. 5F). It is worth emphasizing the need for careful performance comparison between calcium transient detection algorithm, with comparison on open datasets or benchmarks being crucial. Another issue to consider is temporal precision: some early calcium transient detection methods papers reported performance in terms of percentage correct detections and false positives, but with a window for correct detection as long as 0. 5 seconds. We have instead reported detection performance with a window of two sample lengths, or 0.034 sec, a significantly harder criterion to meet (Fig. 5F).

### 6.2 *Analysis of neural population activity*

Having obtained a (possibly large) population of time series (either a continuous time series reflecting calcium or instantaneous firing rate, or a spike train), the next issue is analysis. While this will of course depend upon the specific question being asked in any given study, it is worth considering how the scaling up of dataset dimensionality possible with two photon imaging affects a number of common analyses. One such analysis is receptive field mapping, which (with appropriate stimulus classes, such as white noise) can be performed using spike triggered analysis techniques [162], such as spike triggered average [163], spike triggered covariance [164], and spike triggered independent component analysis [165]. Such techniques, which effectively operate independently on each response element or neuron, apply straightforwardly to two photon imaging data.

**Fig. 5.**
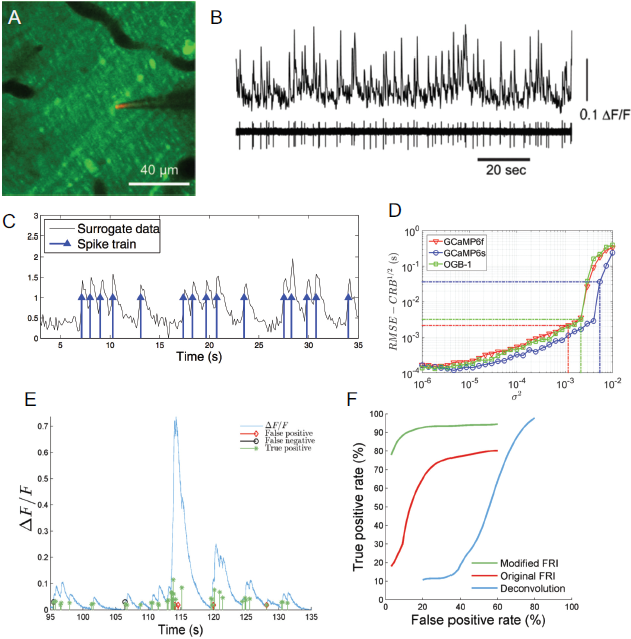
Calcium transient detection. (A) Two photon imaging of calcium signals in cerebellar Purkinje cell dendrites. (B) Simultaneous calcium imaging and electrophysiology (with pipette shown in panel A) to record “ground truth” data. A, B from [158] with permission. (C) Generation of surrogate data. (D) Performance approaches the Cramer-Rao bound with increasing SNR. GCaMP6s provides better detectability than OGB-1 or GCaMP6f. (E) Validation with experimental GCaMP6s data. (F) Receiver Operating Characteristic (ROC) analysis of perfomance on real data, comparing the deconvolution approach of Vogelstein et al. [161] with our original [158] and updated [160] algorithms.

Another common approach is information theoretic analysis [166], in which the question is asked: by how much does knowing a neural response reduce our uncertainty about the value of some external correlate? One way to use this to study how information is represented in the nervous system is to repeat the analysis with varying response codes: if, for instance, one obtains the same amount of information about a sensory stimulus using a pooled code (in which activity is pooled regardless of source) as using a spatial pattern code (in which the activity of each neuron is considered to be a labelled ‘letter’ of a ‘word’), then one can conclude that the pooled code would be the more parsimonious description of the neural coding of that sensory stimulus [see 28, for a counter-example]. An extension of this is to break down the information into components reflecting individual statistical contributions (for instance of single spikes, pairwise correlations in space or time, etc) to the overall information, a procedure which we have termed information component analysis [167, 168]. This approach has been used to dissect out the role of correlations in stimulus encoding [e.g 169]. Related approaches allow the entropy of neural spike patterns (in space or time) to be studied [170]. A major caveat is that information theoretic methods do not scale well with dimensionality [171]. Information theoretic quantities tend to be biased due to the limited number of samples feasible in neurophysiological experiments. While there are increasingly sophisticated estimation algorithms to overcome this [172], such algorithms ultimately cannot overcome the curse of dimensionality, and fail catastrophically as the number of neurons in the pattern becomes too high. One approach to avoiding this problem, and scaling up information theoretic analysis or decoding algorithms, while retaining the ability to analyse pattern contributions is to incorporate models of response probability distributions into the analysis framework [173]. Further advances in the development of algorithms that scale well with pattern size will be important in order to take maximum advantage of novel experimental technologies to image large scale neural circuits.

## 7 Concluding remarks-a roadmap for large-scale imaging of neural circuits

There have been rapid advances in recent years in the development of tools for observing neural circuit function, including novel fluorophores, novel microscopy approaches, and novel data analysis algorithms. These developments have brought their own challenges-from limits on imaging depth due to the scattering of light, to the problems raised by the curse of dimensionality in the analysis of datasets comprised of simultaneous recordings from many neurons. Developments in the near future (see Fig. 6) likely to result in further progress include improved fluorophores with even greater signal amplitude, rapid three dimensional scanning techniques based on devices such as the electrically tunable lens, overcoming scattering limitations through wavefront engineering [174], use of three photon microscopy to take advantage of gaps in the absorption spectra of brain tissue, and the application of lightsheet-like microscopy techniques to intact mammalian preparations, overcoming scanning bottlenecks. Equally as important as these new developments will be the lowering of barriers to the use of these technologies outside of dedicated photonics laboratories-initiatives to make this technology widespread in systems neuroscience laboratories, beyond a few technological astute early adopters, are crucial. While much progress has been made on the development of calcium transient detection algorithms for resolving spike trains in neural circuits, there is a need for robust validation on openly accessible datasets (both real and surrogate). These developments will stimulate the development of algorithms for analysing multi-neuronal activity patterns with improved scaling behaviour, enabling the testing of hypotheses concerning neural circuit function. Furthermore, much can be done directly with continuous time series-whether calcium or voltage based fluorescence signals-and we expect much progress to be made here, with nonlinear dimensionality reduction algorithms for handling the high-dimensional multivariate time series resulting from large-scale optical neural recordings likely to play a crucial role.

**Fig. 6.**
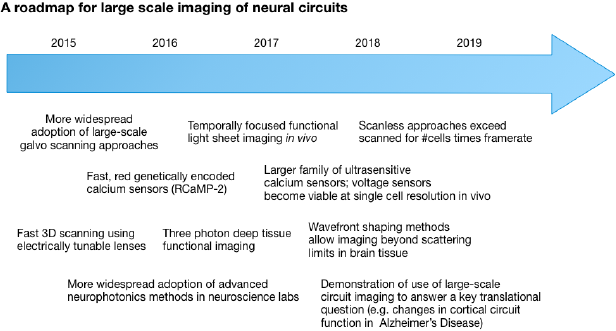
A roadmap for the development of large-scale functional neural circuit imaging technology over the next few years.

